# Neuroprotective pentapeptide, CN-105, improves outcomes in translational models of intracerebral hemorrhage

**DOI:** 10.1101/2020.10.15.339184

**Authors:** Haichen Wang, Timothy D. Faw, Yufeng Lin, Shan Huang, Talaignair N. Venkatraman, Viviana Cantillana, Christopher D. Lascola, Michael L. James, Daniel T. Laskowitz

**Affiliations:** Department of Neurology, Duke University, Durham, NC, USA; Doctor of Physical Therapy Division, Department of Orthopaedic Surgery, Duke University, Durham, NC, USA; Department of Neurology, Tianjin First Central Hospital, Tianjin, China; Department of Geriatrics, Tianjin Medical University General Hospital, Tianjin, China; Department of Radiology, Duke University, Durham, NC, USA; Department of Anesthesiology, Duke University, Durham, NC, USA

**Keywords:** Intracerebral Hemorrhage, CN-105, Apolipoprotein E, Stroke

## Abstract

**Background:** Intracerebral hemorrhage (ICH) is a devastating form of cerebrovascular disease for which there are no approved pharmacological interventions that improve outcomes. Apolipoprotein E (apoE) has emerged as a promising therapeutic target given its neuroprotective properties and ability to modify neuroinflammatory responses. We developed a 5-amino acid peptide, CN-105, that mimics the polar face of the apoE helical domain involved in receptor interactions, readily crosses the blood-brain barrier, and improves outcomes in well-established preclinical ICH models. In the current study, we investigated the therapeutic potential of CN-105 in translational ICH models that account for hypertensive comorbidity, sex, species, and age.

**Methods:** In three separate experiments, we delivered three intravenous doses of CN-105 (up to 0.20 mg/kg) or vehicle to hypertensive male BPH/2J mice, spontaneously hypertensive female rats, or 11-month old male mice within 24-hours of ICH. Neuropathological and neurobehavioral outcomes were determined over 3, 7, and 9 days, respectively.

**Results:** In spontaneously hypertensive male mice, there was a significant dose-dependent effect of CN-105 on vestibulomotor function at 0.05 and 0.20 mg/kg doses (p < 0.05; 95% CI: 0.91 – 153.70 and p < 0.001; 95% CI: 49.54 – 205.62), while 0.20 mg/kg also improved neuroseverity scores (p < 0.05; 95% CI: 0.27 – 11.00) and reduced ipsilateral brain edema (p < 0.05; 95% CI: ^−^0.037 – ^−^0.001). In spontaneously hypertensive female rats, CN-105 (0.05 mg/kg) had a significant effect on vestibulomotor function (p < 0.01; η^2^ = 0.093) and neuroseverity scores (p < 0.05; η^2^ = 0.083), and reduced contralateral edema expansion (p < 0.01; 95% CI: ^−^1.41 – ^−^0.39). In 11-month old male mice, CN-105 had a significant effect on vestibulomotor function (p < 0.001; η^2^ = 0.111) but not neuroseverity scores (p > 0.05; η^2^ = 0.034).

**Conclusions:** Acute treatment with CN-105 improves outcomes in translational ICH models independent of sex, species, age, or hypertensive comorbidity.

## Introduction

Intracerebral hemorrhage (ICH) accounts for approximately 10% of all strokes in the United States. High mortality and poor recovery rates have not demonstrably improved, likely due to lack of therapeutic advancements^1,2^. Therapeutic development might be aided through genetic determinants of functional outcomes as potential targets. Specifically, apolipoprotein (APOE, gene; ApoE, protein) E4 has been associated with poor clinical outcomes following acute brain injury^3^ and increased morbidity and perihematomal edema after ICH^4–6^. Further, association of APOE polymorphism and outcome after ICH has also been replicated in preclinical models^7,8^.

ApoE-mimetic peptides were developed from the apoE receptor binding region to retain the neuroprotective^9^ and anti-inflammatory properties^10^ of the intact protein, while readily penetrating the central nervous system compartment^11^. This approach has demonstrated robust histological and functional improvements in numerous preclinical models of ICH^7,11,12^. Most recently, intravenous administration of the lead candidate apoE-mimetic peptide, CN-105, reduced brain inflammation and development of cerebral edema, while promoting neurobehavioral improvement in a murine model of ICH using adult male mice^11^. Further, CN-105 has received orphan drug status and is currently in early phase clinical trials, including a pilot first-in-disease state trial for patients with primary ICH (NCT03168581, NCT031711903, NCT03802396)^13^. (James et al. 2020, Neurocrit Care; under review)

To improve likelihood of translational success, the present study sought to address additional ICH-relevant paradigms by assessing the efficacy of CN-105 in different species, across sex, and in the setting of major comorbidities (i.e. age, hypertension). We hypothesized that intravenous administration of CN-105 would improve neurobehavioral function and reduce cerebral edema after experimental ICH, independent of species, sex, age, or hypertensive comorbidity.

## Methods

All experiments were approved by and conducted in accordance with the Duke University Institutional Animal Care and Use Committee. Animals were group-housed and maintained on a 12-hour light/dark cycle with ad libitum access to food and water. Animals were randomly assigned into treatment groups prior to injury using a coded study identification number and a computerized randomization protocol with allocation concealment (GraphPad Prism). A block randomization scheme was used so that an equal number of animals were randomized to each of the treatment groups during concurrent experiments. All surgical and nonsurgical procedures were performed in blinded fashion with concealment of treatments, which were coded by an individual blind to group assignment. All *in vivo* experiments were performed to be consistent with the Animal Research Reporting of In Vivo Experiments (ARRIVE) guidelines^14^.

### Experimental Cohorts

As hypertension is the most common comorbidity associated with spontaneous ICH, experiment 1 utilized CN-105 in an ICH model with spontaneously hypertensive male BPH/2J mice (n = 46; 7-11 weeks old; Jackson Labs, Bar Harbor, ME) randomized to Vehicle (n = 11) or CN-105 (0.01 mg/kg, n = 12; 0.05 mg/kg, n = 12; 0.20 mg/kg, n = 11) treatment. Degree of acute cerebral edema (brain water content) and neurobehavioral outcomes were measured over 3 days post-ICH. Next, to extend CN-105 efficacy across species and sex, experiment 2 utilized spontaneously hypertensive, female rats (n = 30; 9-11 weeks old; Envigo, Indianapolis, IN) randomized to Vehicle (n = 15) or CN-105 (0.05 mg/kg, n = 15) treatment. Hematoma volume, cerebral edema progression, and degree of neurobehavioral recovery were measured over 7 days post-ICH. As ICH predominantly occurs in older adults and age has a major impact on outcome, experiment 3 utilized 11-month old male mice (C57BL/6J; n = 25; Jackson Labs, Bar Harbor, ME) randomized into Vehicle (n = 12) or CN-105 (0.05 mg/kg, n = 13) treatment. Neurobehavioral recovery was assessed over 9 days post-ICH. Experimental timelines were strategically selected to investigate impact of comorbid conditions on cerebral edema (Experiments 1 & 2) and acute neurobehavioral recovery (Experiments 2 & 3), while minimizing total animal use. Animals that died following injury but before treatment were excluded from all analyses (Experiment 1: Vehicle, n = 4; CN-105 0.01 mg/kg, n = 3; 0.05 mg/kg, n = 3; 0.20 mg/kg, n = 1; Experiment 2: n = 4 per group; Experiment 3: Vehicle, n = 1; CN-105, n = 2) so that only animals with complete data sets were included (Experiment 1: Vehicle, n = 7; CN-105 0.01 mg/kg, n = 9; 0.05 mg/kg, n = 9; 0.20 mg/kg, n = 10; Experiment 2: n = 11 per group; Experiment 3: n = 11 per group).

### ICH Model

Intrastriatal collagenase injection ocurred as described previously^11,15^. Briefly, rodents were anesthetized with 4.6% isoflurane induction. After tracheal intubation, mechanical ventilation occurred at 0.3 mL of tidal volume and 140 breaths per minute, and anesthesia was maintained with 1.5% isoflurane in a 30% oxygen, 70% nitrogen mixture. We used a rectal probe to monitor body temperature, which was maintained at 37 ± 0.2 °C throughout the procedure via circulating warm water through an underbody pad. A stereotactic frame secured the head during the procedure. We made a midline incision in the scalp to expose the skull and created a burr hole on the left side, lateral to bregma (2.2 mm in mice, 3.2 mm in rats). We delivered type IV-S Clostridial collagenase into the brain over the course of 5 minutes (0.2 U in 0.32 mL normal saline for mice, 1.0 U in 0.40 mL normal saline for rats) via Hamilton syringe (Hamilton Company; Reno, NV) advanced into the cortex (depth = 3 mm in mouse, 5 mm in rat). After collagenase injection, the needle was slowly removed over 5 minutes, and bone wax was used to close the burr hole to avoid back flow. The incision was closed with surgical sutures, and animals were allowed to recover spontaneous respiration and righting reflex prior to extubation.

### Peptide Synthesis and Administration

Polypeptide Inc. (San Diego, CA) synthesized the CN-105 (Ac-VSRRR-amide) peptide to a purity >99% in accordance with good manufacturing practice. The peptide was then dissolved in sterile normal saline, which is the vehicle control for all experiments. For experiment 1, mice received either Vehicle or CN-105 at doses of 0.01 mg/kg, 0.05 mg/kg, or 0.20 mg/kg in saline. For experiments 2 and 3, animals received either Vehicle or CN-105 at 0.05 mg/kg in saline, as this is the lowest effective dose identified from experiment 1 and previous studies^16^. It is also the dose used in early phase clinical trials (NCT03168581, NCT031711903, NCT03802396)^13^. For all experiments, concealed treatment administration occurred at 2, 4, and 24 hours post-ICH via tail vein injection (total volume for each treatment = 100 μL) while animals were gently restrained by a rodent restraining device (Harvard Apparatus; Holliston, MA).

### Rotarod Testing

To determine CN-105 effect on vestibilomotor function, animals underwent testing on an automatically rotating rod (Rotarod; Ugo Basile; Comerio, Italy)^17^. Animals underwent training prior to injury with two consecutive trials of 60 seconds each at 16 rotations per minute. Testing included three trials at accelerating rotational speeds and occurred the day prior to ICH, the day following ICH, and then every other day for up to 9 days post injury. Three test trial latencies were used to calculate average daily rotarod latency for individual animals, which were then used in the statistical analyses. Baseline rotorod latencies were compared prior to injury to ensure there were no between-group differences at the study onset and thus were not not included in post-injury recovery comparisons.

### Neuroseverity Score

We used an observational rating system to evaluate neurobehavioral function prior to ICH and every other day for up to 9 days post injury^7,18^. This system has 7 functional domains that assess spontaneous activity, symmetry, climbing, balance and coordination, body proprioception, vibrissae, and tactile responses. Scores within each domain range from 0 or 1 to 3 (minimum score = 3, maximum = 21), where lower scores denote greater impairment. All animals received the maximum score prior to injury. As such, the average daily neuroseverity scores for each group at each time point post injury were used in the analyses.

### Brain Water Content

To quantify cerebral edema in experiment 1, mice were anesthetized, exsanguinated, and euthanized at 3 days post-ICH. Brains were rapidly dissected and split mid-sagittally into right and left hemispheres on an ice platform. The brainstem and cerebellum were then removed. Each hemisphere was weighed immediately (wet weight) and after dehydrating for 24 hours at 105 °C (dry weight). Cerebral edema was calculated as a percentage of wet weight (percent water content = [wet weight – dry weight] / [wet weight] x 100) and expressed as percent water content.

### Magnetic Resonance Imaging (MRI) Analysis of Hemorrhage Volume and Cerebral Edema

Imaging was performed using a Bruker 7.0T scanner (Bruker Biospin, Billerica, MA, USA) operating with Paravision 5.1. Animals were anesthetized with 1.5% isoflurane in room air. Core body temperature was maintained at 37 ± 0.5 °C. Fast spin echo-based T 2 -weighted acquisitions were performed using echo time (TE)/repetition time (TR) = 12/4,200 ms, field of view (FOV) = 3 × 3 cm, matrix 128 × 128, 32 slices, and 1.0 mm slice thickness. Susceptibility Weighted Images (SWI) were collected using a FLASH sequence with the following parameters: TE/TR = 9/700 ms, flip angle (FA) = 40, FOV = 3 cm x 3 cm, matrix size = 256 × 256, thickness = 0.5 mm, slices = 32. Diffusion-weighted images (DWI) and apparent diffusion coefficient (ADC) maps were generated using double spin-echo planar pulse sequence usingthree orthogonal directions. Four b-values (b = 0, 100, 200, 1000) were acquired with a matrix size of 128 × 128 × 15, slice thickness 1.0 mm; total scan time = 27 minutes. SWI images were reconstructed using Paravison. Hemorrhage volumes and brain free water were measured off-line in Osirix using manually selected ROIs at 2 and 24 hours after injury. As the T_2_ signal is a function of intracellular and extracellular water and susceptibility, measuring T_2_ in the contralateral hemisphere, distant from the site of hemorrhage, avoids the contribution of susceptibility from blood products and is reflective of global rather than localized changes in edema. Moreover, concomitant DWI confirms that there should be no added T_2_ signal contribution from intracellular water in this region, which would correspond to cell swelling or cytotoxic edema. Thus, T_2_ changes in this contralateral region should reflect changes in extracellular edema alone. To assess progression of cerebral edema after injury, T_2_-weighted images from the contralateral frontal cortex were normalized to precranial T_2_ signal in muscles of mastication, and these normalized T_2_ values (T_2_ frontal lobe / T_2_ muscle) were compared at 2 and 24 hours post injury. Positive values indicate increased brain free water over time.

### Statistical Analyses

Functional outcomes (rotarod latency and neuroseverity score) were analyzed using two-way analysis of variance (ANOVA). Analysis of neuropathological outcomes (brain water content, hemorrhage volume, contralateral edema expansion) occurred using either one-way ANOVA (experiment 1) or independent samples t-test (experiment 2). Where appropriate, Dunnett’s *post hoc* test was used to determine between-group differences. Means and standard error of the mean (SEM) are reported throughout. Significance was set at p < 0.05 for all statistical tests.

## Results

### CN-105 improves neurobehavioral function and reduces cerebral edema in hypertensive, male mice

To determine whether CN-105 improves recovery after ICH in the setting of comorbid hypertension, we induced ICH in spontaneously hypertensive, male BPH/2J mice and assessed neurobehavioral function over 3 days. There was no difference in rotarod latency between groups (Vehicle = 242.1 ± 9.4 sec; CN-105, 0.01 mg/kg = 232.3 ± 8.7 sec, 0.05 mg/kg = 235.9 ± 9.0 sec, 0.20 mg/kg = 250.7 ± 8.8 sec; p > 0.05) and all mice received the maximum neuroseverity score (21) prior to ICH induction. There was a significant effect of treatment on vestibulomotor function in hypertensive mice (Fs = 6.211, DF = 3, p < 0.001). Treatment with CN-105 at either 0.05 or 0.20 mg/kg signifncantly improved vestibulomotor functional recovery compared to Vehicle (p = 0.047 and p < 0.001, respectively; Dunett’s *post hoc*; Fig. 1A). A one-way ANOVA revealed a trend of improved neuroseverity scores at 3 days post injury with CN-105 treatment (Fs = 2.235, DF = 3; p = 0.098). *Post hoc* testing showed that hypertensive mice treated with 0.20 mg/kg CN-105 had significantly higher neuroseverity scores at 3 days post-ICH (p = 0.037; Dunnett’s *post hoc*; Fig. 1B).

**Fig. 1.**
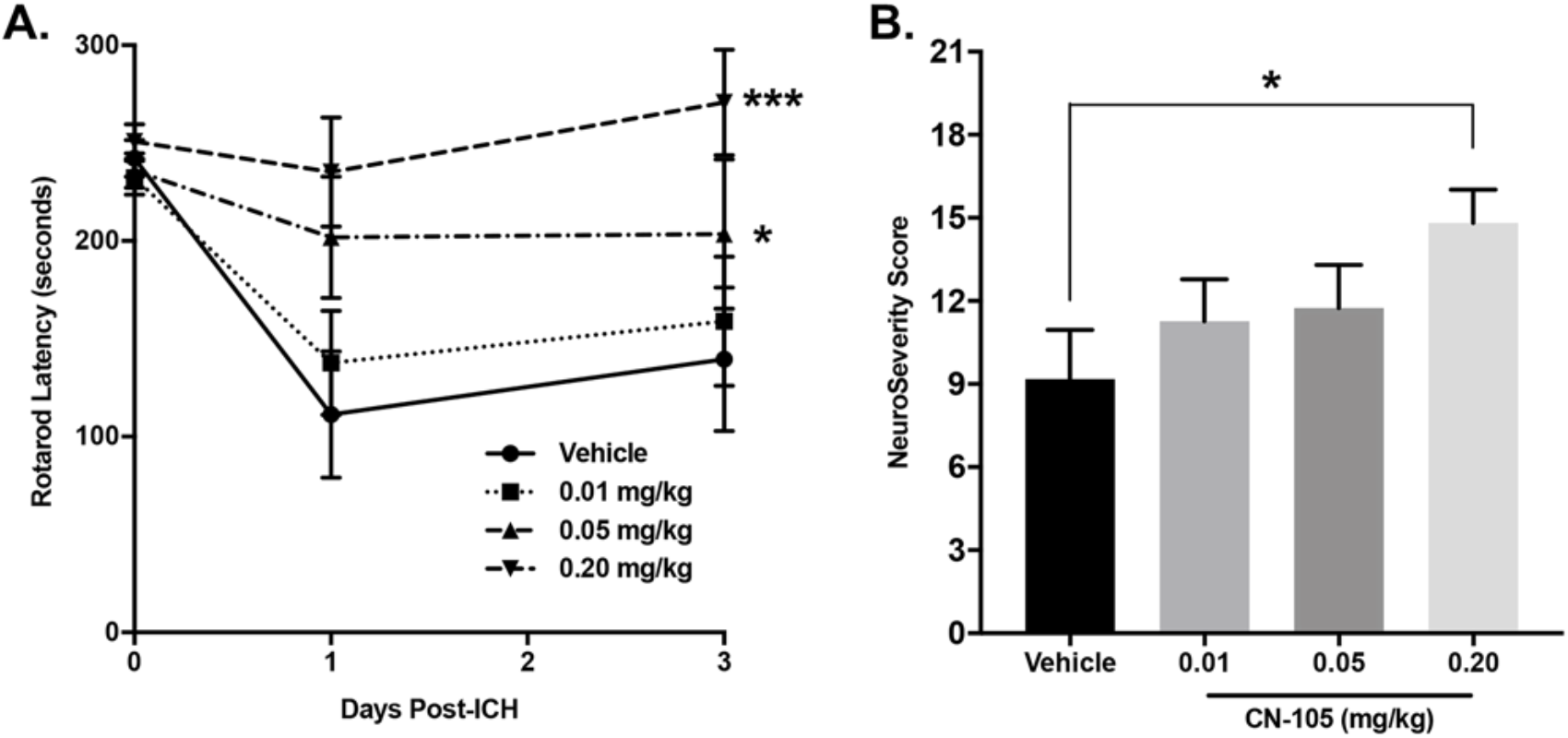
CN-105 improves neurobehavioral function in hypertensive, male mice with ICH. Spontaneously hypertensive BPH/2J mice with ICH received Vehicle (n = 7) or CN-105 treatment (0.01 mg/kg, n = 9; 0.05 mg/kg, n = 9; 0.20 mg/kg, n = 10). A) Vestibulomotor function was assessed prior to injury, 1 and 3 days after injury via rotarod. CN-105 treatment at 0.05 mg/kg and 0.20 mg/kg had a significant effect on recovery of vestibilomotor function represented by rotarod latency (***p < 0.001; Two-way ANOVA with Dunnett’s *Post Hoc*). B) Neuroseverity scores were assessed on day 3 post injury. Treatment with CN-105 at 0.2 mg/kg reduced neurological deficits based on neuroseverity scoring (*p < 0.05; One-way ANOVA with Dunnett’s *Post Hoc*).

Next, we determined the effect of CN-105 treatment on ICH-induced brain edema in hypertensive male mice. There was a trend toward an overall main effect of CN-105 treatment that did not reach statistical significance (Fs = 2.51, DF = 3; p = 0.079). *Post hoc* comparisons revealed that ipsilesional brain water content was significantly reduced in mice treated with 0.20 mg/kg CN-105 compared to Vehicle (p = 0.038; Dunnett’s *post hoc*; Fig. 2). These data indicate that treatment with 0.20 mg/kg CN-105 improves recovery and reduces brain edema after ICH in hypertensive male mice.

**Fig. 2.**
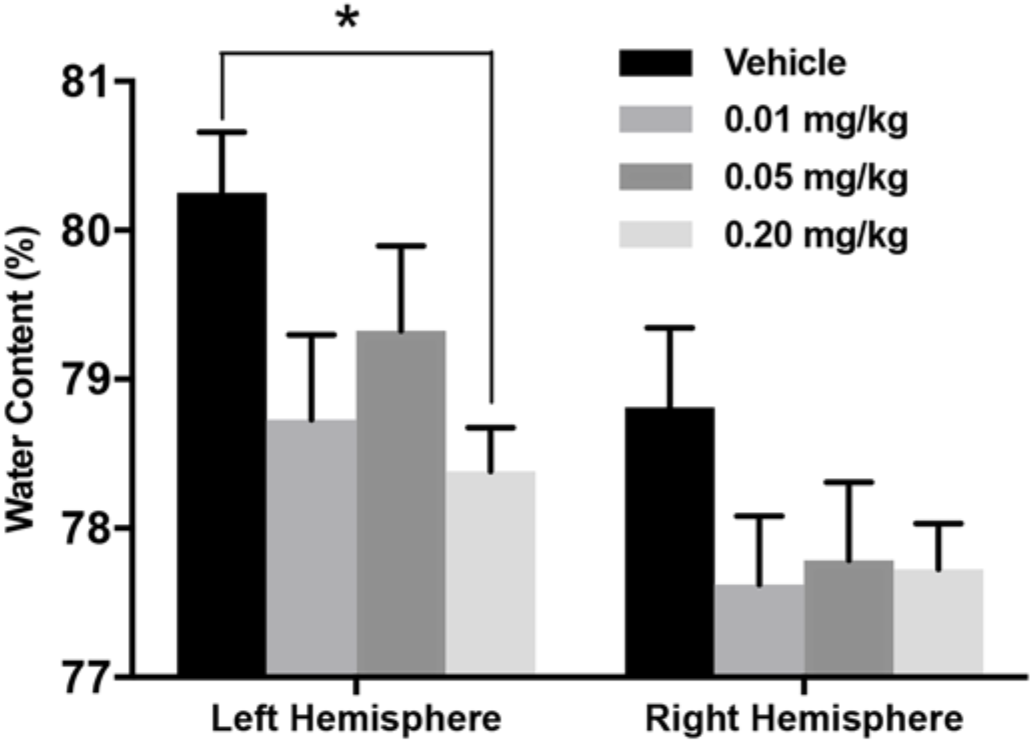
CN-105 reduces cerebral edema in hypertensive, male mice with ICH. Spontaneously hypertensive BPH/2J mice with ICH were treated with Vehicle (n = 7) or CN-105 (0.01 mg/kg, n = 9; 0.05 mg/kg, n = 9; 0.20 mg/kg, n = 10) and cerebral edema was assessed at 3 days post injury. Ipsilateral hemisphere water content was reduced in mice treated with 0.20 mg/kg CN-105 (*p < 0.05; One-way ANOVA with Dunnett’s *Post Hoc*). Abbreviations: ICH, Intracerebral Hemorrhage

### CN-105 improves neurobehavioral function and reduces progression of cerebral edema in spontaneously hypertensive female rats

To determine the effectiveness of CN-105 in a second species and sex with hypertensive comorbidity, we induced ICH in spontaneously hypertensive, female rats and assessed functional recovery every other day for the first week post injury. Again, there was no difference in rotatod latency (Vehicle = 370.4 ± 30.7 sec; CN-105 = 356.7 ± 16.7 sec; p > 0.05) or neuroseverity score (all animals = 21) prior to injury. There were significant effects of CN-105 treatment on both latency to fall (Fs = 8.16, DF = 1; p = 0.005; Fig 3A) and neuroseverity scores (Fs = 6.88, DF = 1; p = 0.011; Fig 3B).

**Fig. 3.**
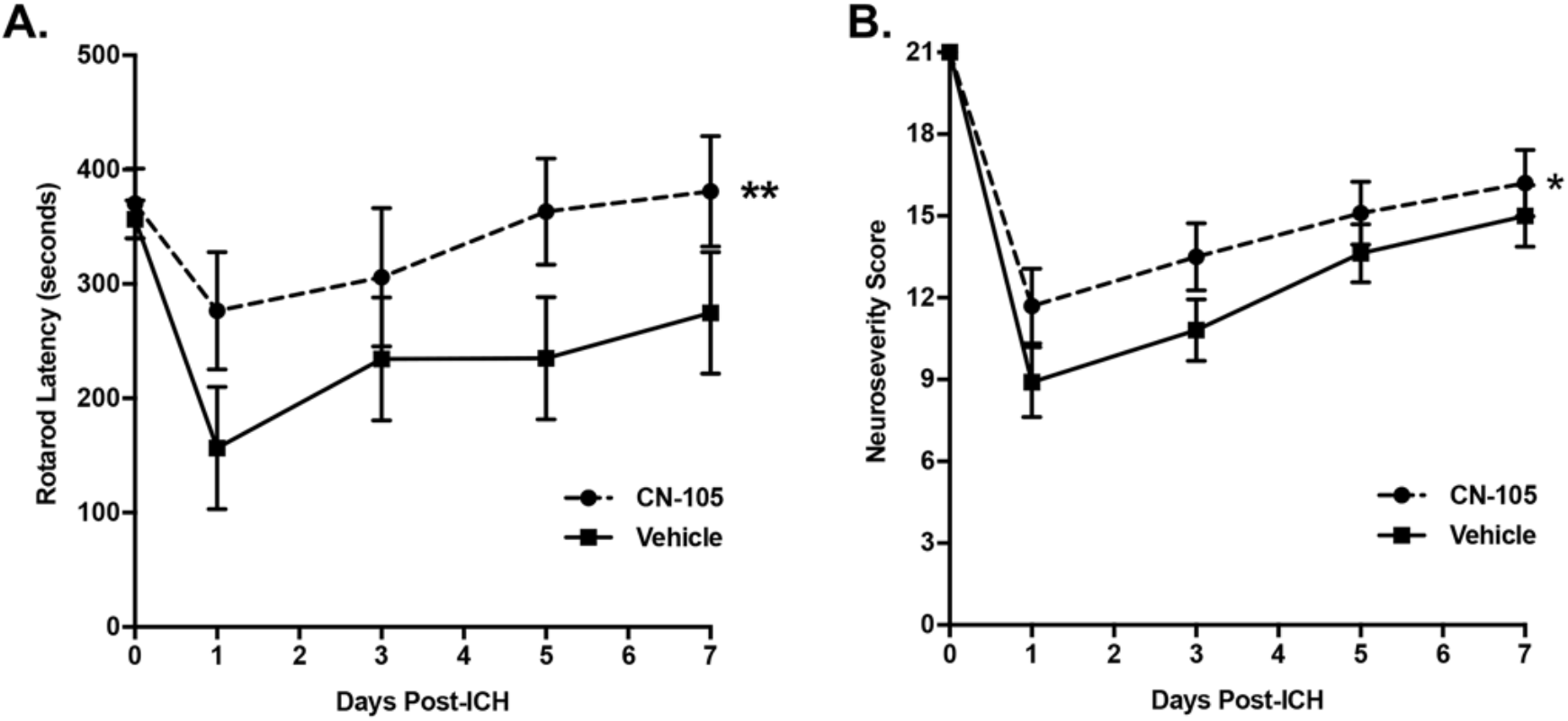
CN-105 improves neurobehavioral function in spontaneously hypertensive female rats. Rats with ICH received either Vehicle or CN-105 treatment (n = 11 per group) and neurologic function was assessed prior to injury and daily for 7 days post injury. Both rotarod latency and neuroseverity score increased in CN-105 treated group over 7 days following ICH. CN-105 robustly improved rotarod performance (A, **p < 0.01; Two-way ANOVA). CN-105 also significantly improved neurological scores (B, *p < 0.05; Two-way ANOVA). Abbreviations: ICH, Intracerebral Hemorrhage

We next determined the effect of CN-105 treatment on hemorrhage volume (Fig 4A,B) and edema progression (Fig 4C,D) in hypertensive female rats with ICH. Similar to our previous results^11^, CN-105 treatment did not reduce hemorrhage volume at 2 or 48 hours post-ICH (p > 0.05; Fig 4B). However, rats treated with CN-105 demonstrated significantly less edema progression in the contralateral hemisphere measured by MRI (p = 0.005; Fig 4B).

**Fig. 4.**
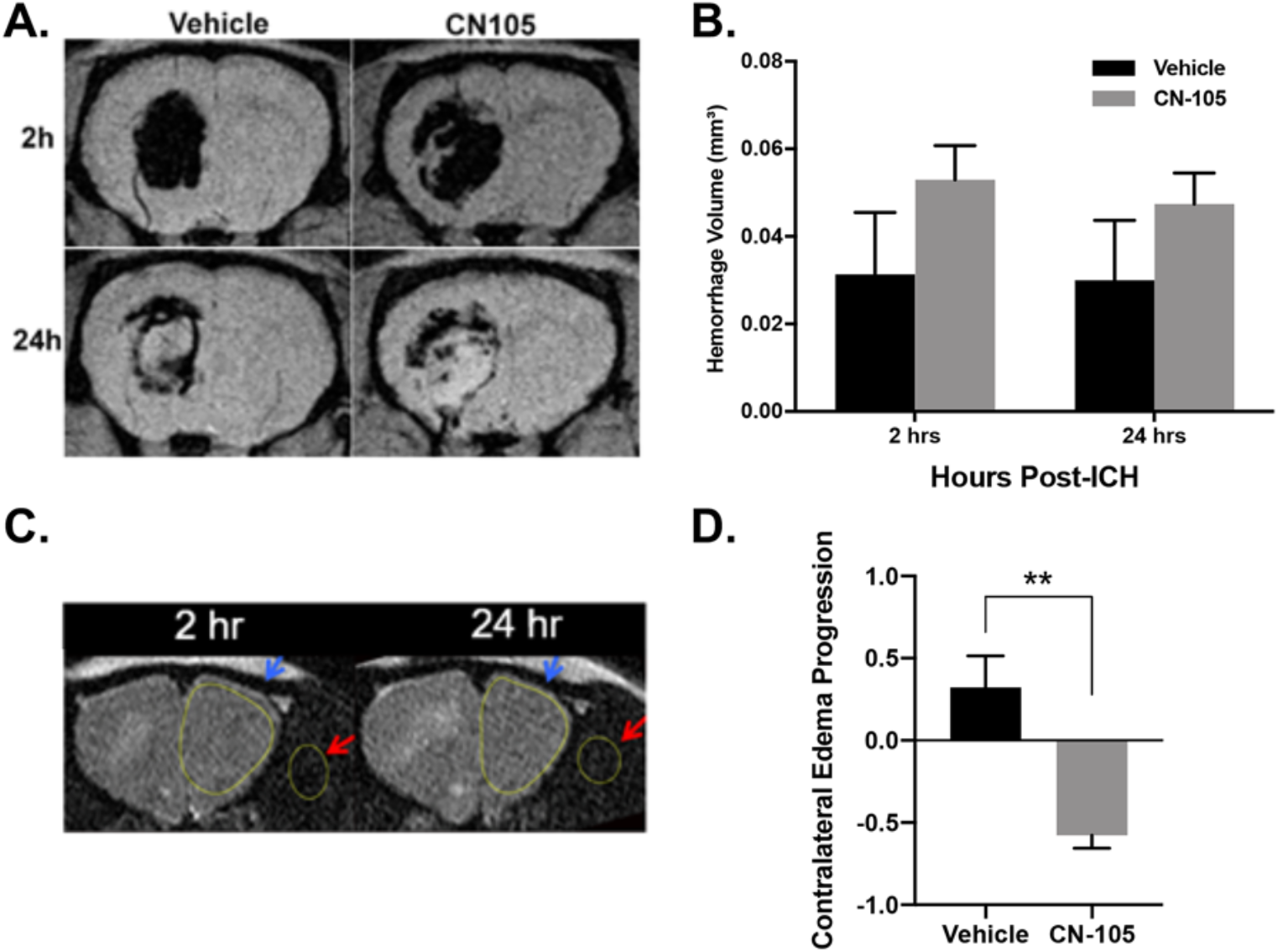
CN-105 reduces progression of cerebral edema in spontaneously hypertensive female rats. Rats with ICH received either Vehicle or CN-105 treatment (n = 11 per group). A) Hemorrhage volume was assessed 2 and 24 hours post-ICH using MRI. B) There were no differences in hemorrhage volume between Vehicle and CN-105 groups at either timepoint (NS, p > 0.05; Independent Samples t-Test). C) Representative images at 2 and 24 hours post-ICH illustrate contralateral frontal lobe (blue arrows) and extra-cranial muscles (red arrows) utilized to quantify edema progression. D) Progression of contralateral brain free water was significantly decreased with CN-105 treatment indicating reduced evolution of cerebral edema (**p = 0.0049; Independent Samples t-Test). Abbreviations: ICH, Intracerebral Hemorrhage; MRI, Magnetic Resonance Imaging

### CN-105 improves neurobehavioral function in middle-aged, male mice

Age is a major predictor of ICH outcome in the clinical setting. As such, we sought to determine the effectiveness of CN-105 in 11-month old mice after ICH. Rotarod latencies (Vehicle = 233.3 ± 20.0 sec; CN-105 = 211.7 ± 15.8 sec; p > 0.05) and neuroseverity scores (all animals = 21) were similar between groups prior to injury. After ICH, rotarod performance was significantly different between mice treated with CN-105 compared to Vehicle treatment (Fs = 12.44, DF = 1; p < 0.001; Fig 5A). There was a trend toward improved neuroseverity scores in the older mice treated with CN-105, but this did not reach statistical significance (Fs = 3.53, DF = 1; p = 0.063; Fig 5B).

**Fig. 51.**
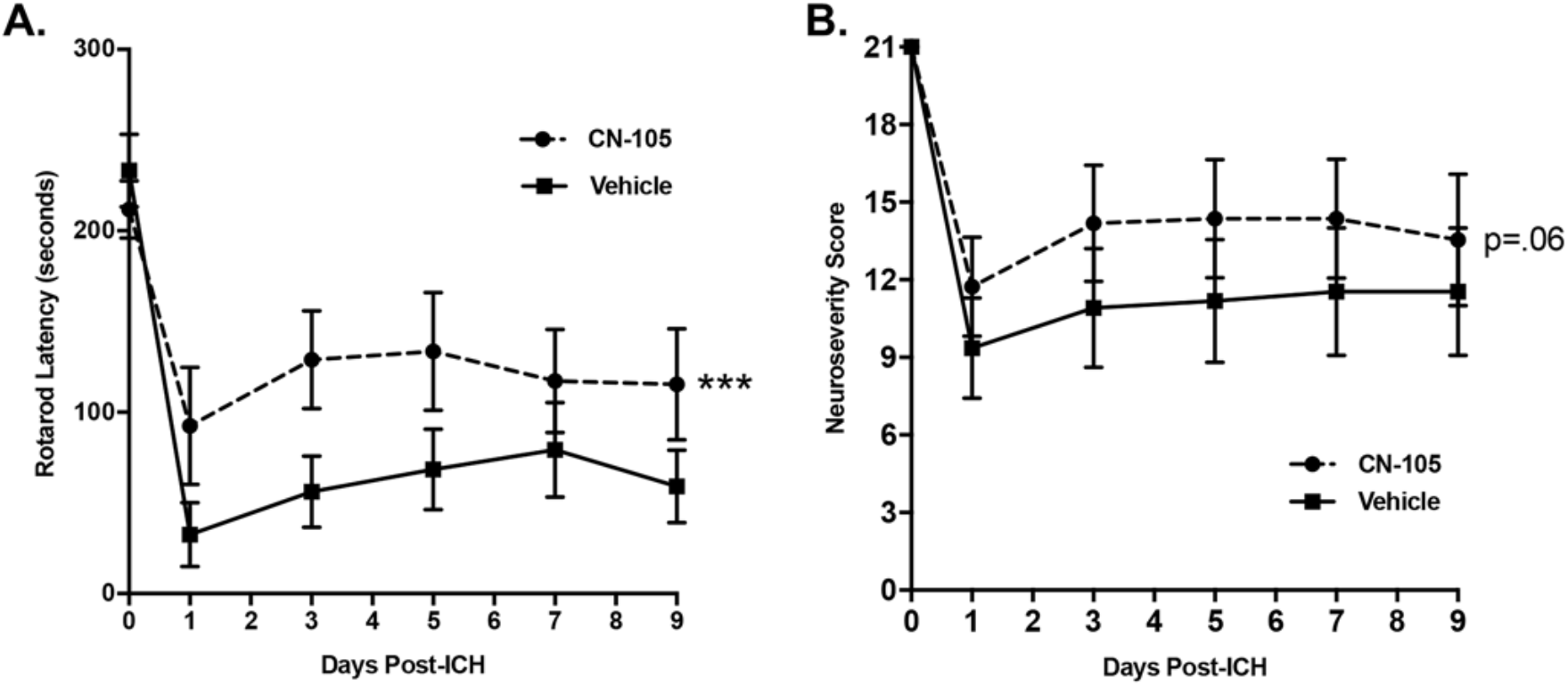
CN-105 improves neurobehavioral function in middle-aged, male mice. Eleven-month old mice with ICH received Vehicle or CN-105 treatment (n = 11 per group). Neurobehavioral testing occurred prior to injury and every other day for 9 days post injury. A) CN-105 treatment had a significant effect on vestibulomotor function (***p < 0.001; Two-way ANOVA). B) The trend toward improved performance associated with CN-105 treatment did not reach statistical significance for the neuroseverity scores (NS, p = 0.06; Two-way ANOVA) in 11-month old mice. Abbreviations: ICH, Intracerebral Hemorrhage

## Discussion

The inability of promising preclinical therapeutic candidates to translate to clinical efficacy has led to a critical unmet need for effective treatments to reduce the devastating effects of acute brain injury^19^. According to the Hemorrhagic Stroke Academia Industry (HEADS and HEADS-2) roundtable recommendations^20,21^, to improve the likelihood of successful translation, preclinical research of therapeutic candidates for ICH should include hypertensive models, demonstrate efficacy on meaningful neurological deficits, provide evidence that the therapeutic candidate does not increase bleeding risk or hematoma expansion, demonstrate evidence for safety in cerebral ischemia, demonstrate replicability, and show evidence of efficacy across species, sex, age, and ICH models. Moreover, the therapeutic time window should be realistic for human use with evidence that drug reaches the therapeutic target.

Acute brain injury responses are modified by apoE, a 299-amino acid protein, produced within the brain. Data from our lab and others demonstrate that apoE exerts neuroprotective effects by downregulating neuroinflammatory responses^22,23^ and that these biological activities can be replicated in smaller apoE-mimetic peptides^10^. The inability of exogenous apoE holoprotein to cross the blood-brain barrier led to the development of the 5-amino acid peptide, CN-105, derived from the polar face of the receptor binding region. We previously demonstrated that CN-105 improves short- and long-term neurobehavioral and histological outcomes, decreases cerebral edema, and increases neuronal survival after ICH in adult male mice^11^. Here, we extend those findings across key, clinically-relevant translational ICH models. These data provide the first evidence of improved functional and neuropathological outcomes associated with CN-105 treatment in spontaneously hypertensive male mice, spontaneously hypertensive female rats, and middle-aged male mice, meeting many of the established guidelines above to increase successful translational likelihood. That CN-105 yields postitive outcomes after ICH, independent of animal strain, species, sex, and comorbid conditions, provides strong support for further testing of CN-105 in diverse clinical ICH populations.

Hypertension is the most significant risk factor for non-lobar ICH and is present in up to 80% of all patients who suffer ICH^24^. Moreover, hypertension plays a major role in post-ICH hematoma expansion, cerebral edema, and secondary neuronal injury^25,26^. Early phase clinical trials are not adequately powered to rigorously evaluate potential interactions between therapeutic effect and hypertension; thus, preclinical studies should be performed in the presence of comorbid hypertension to facilitate translation of promising therapies. For example, while more than 50% of patients enrolled into the early phase safety and feasibility trial (NCT031711903) were hypertensive, sample size prohibits definitive determination of interaction of hypertension on relevant clinical outcomes in the setting of CN-105 treatment after acute ICH. Here, we utilized two preclinical models of spontaneous hypertension to test the efficacy of CN-105. In both spontaneously hypertensive male BPH/2J mice and spontaneously hypertensive female rats, administration of CN-105 improved neurobehavioral outcomes and reduced cerebral edema, suggesting efficacy in the setting of this ICH-relevant comorbidity. Further, reduction of cerebral edema evolution in spontaneously hypertensive rats, as measured by MRI, was observed in the hemisphere contralateral to focal ICH. This is consistent with our observations that even focal injuries may cause diffuse microgliosis and involvement of the contralateral hemisphere^27^. Although contralataeral edema is often not appreciated in clinical imaging studies, long-term cognitive deficits after focal brain injuries in humans may be a result of more diffuse brain involvement. Present results also provide additional evidence supporting the role of CN-105 in limiting secondary effects of ICH.

Age is another major outcome determinant for spontaneous hypertensive ICH. Notably, 11-month old mice used here are representative of middle-aged rather than a geriatric population, which is consistent with the average age of individuals who experience ICH^2,28,29^. A number of reports have suggested that the risk of in-hospital mortality increases with advancing age^30^. Two proposed mechanisms for age-associated negative effects on ICH outcomes include increased hematoma volume^31,32^ and neuroinflammation^32,33^. Previous reports show hematoma volume is not affected by APOE genotype^6^ or exogenous administration of apoE-mimetic therapies^11^. Consistent with these findings, hematoma volume was unaffected by treatment with CN-105 across the present set of experiments. However, we have repeatedly shown beneficial effects of CN-105 treatment on neuroinflammation through reduction of glial activation in various models of acute brain injury^11,16,27,34^. Given that microglia are a major source of the hyperinflammatory response observed in aging^35,36^, the ability of CN-105 to reduce glial inflammatory responses may prove especially beneficial for patients with increased age.

There are several limitations to our current findings. Although injection of clostridial collagenase into the basal ganglia recapitulates many of the cardinal features of deep ICH, including small vessel rupture and hematoma evolution, it is less representative of the pathophysiology of lobar hemorrhages. Given known differences in lobar and non-lobar hemorrhage^37,38^, other complementary models might also be tested for therapeutic effect of CN-105, namely, autologous blood injection^15^. Of course, early phase trials may demonstrate efficacy in one ICH sub-type (hypertensive; non-lobar) over another (amyloid angiopathy; lobar) obviating the need for testing in other preclinical models. Additionally, while we demonstrate efficacy in both mouse and rat, these represent lissencephalic species, and efficacy in a gyrencephalic model might also add confidence for CN-105’s translational potential. However, preliminary data suggest CN-105 improved functional outcomes in a (gyrencephalic) ferret blast injury model as well^39^. Lastly, while CN-105 was rationally developed from the polar face of the apoE receptor binding region and overwhelming preclinical evidence establishes its neuroprotective properties^40^ and that its anti-inflammatory effects are mediated via the LRP-1 receptor^41–44^, CN-105’s mechanism of action after acute ICH has not been definitively established.

In conclusion, we found that CN-105, a 5-amino acid peptide derived from the polar face of the apoE receptor binding region reduces cerebral edema, and improves functional outcomes in clinically-relevant experimental ICH paradigms induced by collagenase injection. The beneficial effects are consistently found independent of species, sex, or the presence of comorbid hypertension. Given the favorable pharmacokinetic profile and safety signal in early Phase 1^13^ and disease-specific trials in acute ICH (NCT03168581, NCT03711903), CN-105 represents an attractive candidate for clinical translation.

## Data Availability

The data that support the findings of this study are available from the corresponding author upon reasonable request.

## Author Contributions

HW, MLJ, and DTL designed the research. HW, YL, SH, VC, and TNV performed the research and collected the data. HW, DTL, CDL, VC, TNV, MLJ, and TDF analyzed and interpreted the data. TDF, HW, MLJ, and DTL wrote the manuscript. HW, TDF, and CDL prepared the figures. All authors edited the manuscript and approved the final version.

